# Expression of ACE2, TMPRSS2 and CTSL in human airway epithelial cells under physiological and pathological conditions: Implications for SARS-CoV-2 infection

**DOI:** 10.1101/2020.08.06.240796

**Authors:** Junping Yin, Brigitte Kasper, Frank Petersen, Xinhua Yu

## Abstract

SARS-CoV-2 enters into human airway epithelial cells via membrane fusion or endocytosis, and this process is dependent on ACE2, TMPRSS2, and cathepsin L. In this study, we examined the expression profiles of the three SARS-CoV-2 entry-related genes in primary human airway epithelial cells isolated from donors with different physiological and pathological backgrounds such as smoking, COPD, asthma, lung cancer, allergic rhinitis, cystic fibrosis, or viral infections. By reanalyzing 54 GEO datasets comprising transcriptomic data of 3428 samples, this study revealed that i) smoking is associated with an increased expression of ACE2 and TMPRSS2 and a decreased expression of cathepsin L; ii) infection of rhinovirus as well as poly(I:C) stimulation leads to high expression of all three SARS-CoV-2 entry-related genes; iii) expression of ACE2 and cathepsin L in nasal epithelial cells are decreased in patients with asthma and allergic rhinitis. In conclusion, this study implicates that infection of respiratory viruses, cigarette smoking and allergic respiratory diseases might affect the susceptibility to and the development of COVID-19.

## Introduction

In December 2019, a cluster of viral pneumonia cases which is featured by pulmonary parenchymal opacities at chest radiography was reported in Wuhan, China ^1^. Analysis of lower respiratory tract samples from patients identified a novel beta-coronavirus termed severe acute respiratory syndrome coronavirus 2 (SARS-CoV-2) as the casual pathogen for the pneumonia ^2^. On 11 March 2020, the World Health Organization (WHO) declared coronavirus disease 2019 (COVID-19) as a pandemic ^3^. As of June 29, 2020, there have been over ten million laboratory-confirmed cases of COVID19 worldwide with more than a half million deaths ^4^. SARS-CoV-2 shares approximately 80% sequence identify to SARS-CoV and both viruses use the same cell entry receptor, namely *angiotensin converting enzyme 2 (*ACE2) ^5, 6^. As the receptor of SARS-CoV and SARS-CoV-2, ACE2 mediates the viral entry via two major pathways, cathpesine L-dependent endocytosis and transmembrane serine protease 2 (TMPRSS2) dependent membrane fusion ^7–9^. Since COVID-19 is an acute disease resulted from respiratory tract infection of SARS-CoV-2, the interaction between the spike protein (S protein) of the virus and the ACE2 on human airway epithelial cells is a crucial step for the development of the disease.

Although SARS-CoV-2 can infect individuals of any age, most of the severe cases have been reported in older adults or patients with significant comorbidities including chronic respiratory diseases ^10^. For example, it has been demonstrated that Chronic Obstructive Pulmonary Disease (COPD) is associated with a significant, roughly five-fold increased risk for the development of severe COVID-19 infections ^11, 12^. Understanding the molecular mechanisms behind the increased risk for severe COVID-19 in patients with chronic respiratory diseases would provide clues toward their pathophysiology and the identification of *therapeutic* targets. Since ACE2 might play an important role in protecting the host against lung injury ^13^, one hypothesis is that the expression of ACE2 in airway epithelial cells of patients with chronic respiratory diseases might be upregulated, which enhances the infection of SARS-CoV-2. This notion is partially supported by a recent study, in which Leung et al. reported that smokers and patients with COPD show a higher levels ACE-2 than healthy non-smokers ^14^.

Here, our objective is to draw a complete picture of the expression of three genes essentially involved in SARS-CoV-2 entry, namely ACE2, TMPRSS2, and cathepsin L in human airway epithelial cells under both physiological and pathological conditions. To reach this aim, we retrieved previously published transcriptomic datasets of human airway epithelial cells of healthy subjects with different status of cigarette smoking as well as patients with various respiratory diseases, including COPD, asthma, allergic rhinitis, lung cancer, cystic fibrosis and viral infection.

## Results

### Study selection and data retrieval

A search of the Gene Expression Omnibus (GEO) database with the key word “Airway epithelial cells” resulted in 5065 hits. In the next step, 642 non-human items, 4099 non-series items, and 153 items of non-gene expression array were filtered out. Subsequent assessment further excluded 117 items which were transcriptomic data of cell lines, redundant data, or datasets containing only one group or less than 5 samples per group. Finally, 54 GEO datasets comprising 3428 transcriptomic data of human airway epithelial cells were identified (Fig. 1, Supplementary Fig. 1). Microarray data of each dataset were retrieved from the GEO database. After combining datasets generated by same research group with identical platforms, 28 transcriptomic datasets were generated and used for further analysis (Supplementary table 1). After background correction and normalization of each transcriptomic dataset, expression levels of ACE2, TMPRSS2 and cathepsin L were used for further analysis.

**Fig. 1.**
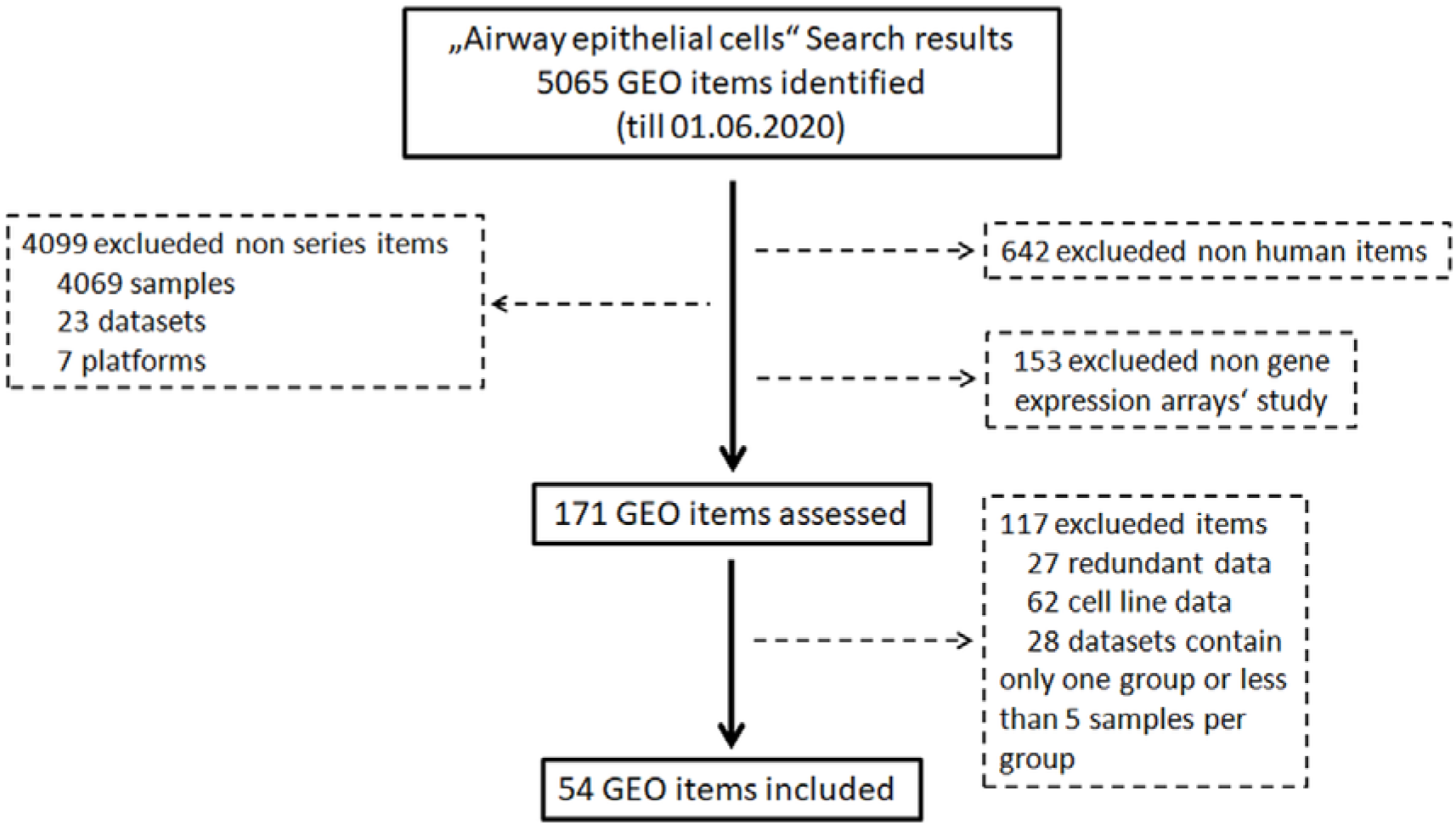
Flow chart of GEO databases screening and selection process.

### Cigarette smoking affects expression of ACE2, TMPRSS2 and cathepsin L in human airway epithelial cells

To examine the expression of the three genes in airway epithelial cells under physiological conditions, we performed comparisons among healthy individuals according to their status of cigarette smoking. The first comparison was performed between current smokers and never smokers. In total, 12 comparisons were performed, with 594 current smokers and 329 never smokers ^15–31^. As shown in Fig. 2a, mean values of expression levels of AEC2 in current smoker were consistently higher than those in never smokers in all 12 comparisons, and differences in 6 out of 12 were statistically significant. Meta-analysis for the standardised mean difference (SMD) showed a moderate effect of smoking on ACE2 expression (SMD=0.70, P<0.001) (Fig. 2b). Similar to ACE2 expression, the expression of TMPRSS2 was also significantly increased in airway epithelial cells of current smoker as compared to never smokers (SMD=0.64, P=0.001) (Fig.2 a,b). By contrast, smoking affected the expression of cathepsin L in the opposite direction. As compared to never smokers, current smokers expressed significantly lower levels of cathepsin L (SMD=−0.43, P<0.001) (Fig. 2a,b).

**Fig. 2.**
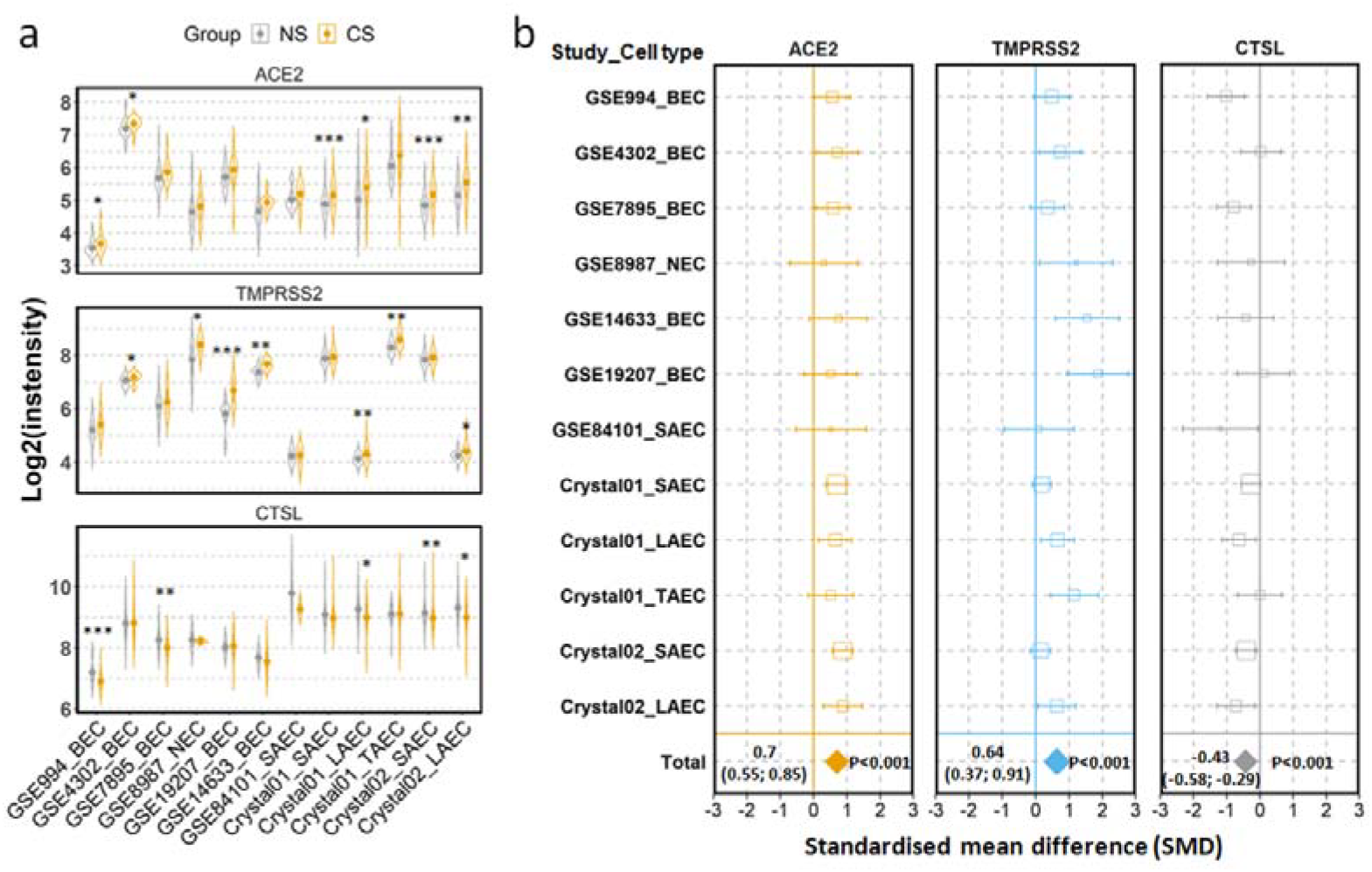
Expression of ACE2, TMPRSS2 and cathepsin L (CTSL) in airway epithelial cells of healthy current smokers (CS) and never smokers (NS). **a)** Violin plot of expression levels of ACE2, TMPRSS2 and CTSL in 12 datasets which contain current smokers and never smoker. Mean and standard deviation (SD) of each group are presented as dot and line, respectively. Statistical difference was calculated by Student’s t test. *p<0.05, **p<0.01, ***p<0.001. **b)** Forest plot of 12 datasets examining expression of ACE2, TMPRSS2 and CTSL in current smokers and never smokers. The x-axis indicates the standardized mean difference (SMD), while the y-axis shows GEO datasets and cell types. Each square in the plots represents the SMD in corresponding datasets and the 95% confidence interval (CI) is shown by the error bar. The size of each square represents the weight of the individual dataset in the meta-analysis. The diamonds in the bottom represent the SMD of the meta-analysis. The SMD, 95% CI and *P* values of meta-analysis are depicted. BEC, bronchial epithelial cell, NEC, nasal epithelial cell, SAEC, small airway epithelial cell, LAEC, large airway epithelial cell, TAEC, trachea airway epithelial cell.

We next investigated whether the effect of cigarette smoking on the three genes reflects a chronic or an acute response of the tissue. Six datasets which contain expression data of former smokers and current smokers were recruited for this comparison ^16, 23, 29, 32–34^. Notably, the differences in the expression of the three genes between current smokers and former smokers follow the same pattern observed between current smokers and never smokers. As compared to former smokers, current smokers showed a moderate increased expression levels of ACE2 and TMPRSS2, which was found to be significant only for the latter protein (ACE2;SMD=0.58, *P*=0.11 and TMPRSS2; SMD=0.27, P=0.003, respectively). By contrast, decreased levels were seen for cathepsin L (SMD=−0.31, P=0.046) (Supplementary Fig. 2). In line with this observation, a comparison between never smokers and former smokers ^16, 23, 29^ revealed no significant differences between these group with regard to the expression of this genes (Supplementary Fig. 3). Thus, these results suggest that the effect of cigarette smoking on expression of ACE2, TMPRSS2 and cathepsin L is more likely an acute reaction than a chronic alteration of the tissue.

The acute effect of cigarette smoking on ACE2 was confirmed in a further dataset ^35^. In this study, current smokers were asked to refrain from cigarette smoking for at least 2 days and then subjected to acute smoking exposure. As shown in Supplementary Fig. 4, the acute smoking exposure increased the expression of ACE2, but showed no effect on TMPRSS2 or cathepsin L.

### Expression of ACE2, TMPRSS2 and cathepsin L in COPD, lung cancer and cystic fibrosis

Consequently, we compared in the next step expression profiles of the three SARS-CoV-2 entry-related genes of epithelial cells derived from none-diseased subjects with those of patients with different respiratory disorders. Since smoking has a major impact on the expression of these genes, we focused on COPD as a smoking-related disease in a first approach.

Three datasets were recruited to compare never smoker control and COPD patients ^15, 17–21, 25, 26, 28, 31, 36^. Expression of ACE2 and TMPRSS2 were found to be increased, while the expression of cathepsin L was decreased in COPD patients as compared to never smoker controls (Fig. 3a-b). Meta-analysis revealed a strong effect of COPD on ACE2 expression (SMD=0.82, P<0.0001), a moderate effect on TMPRSS2 (SMD=0.57, P<0.0001), and no significant effect on cathepsin L.

**Fig. 3.**
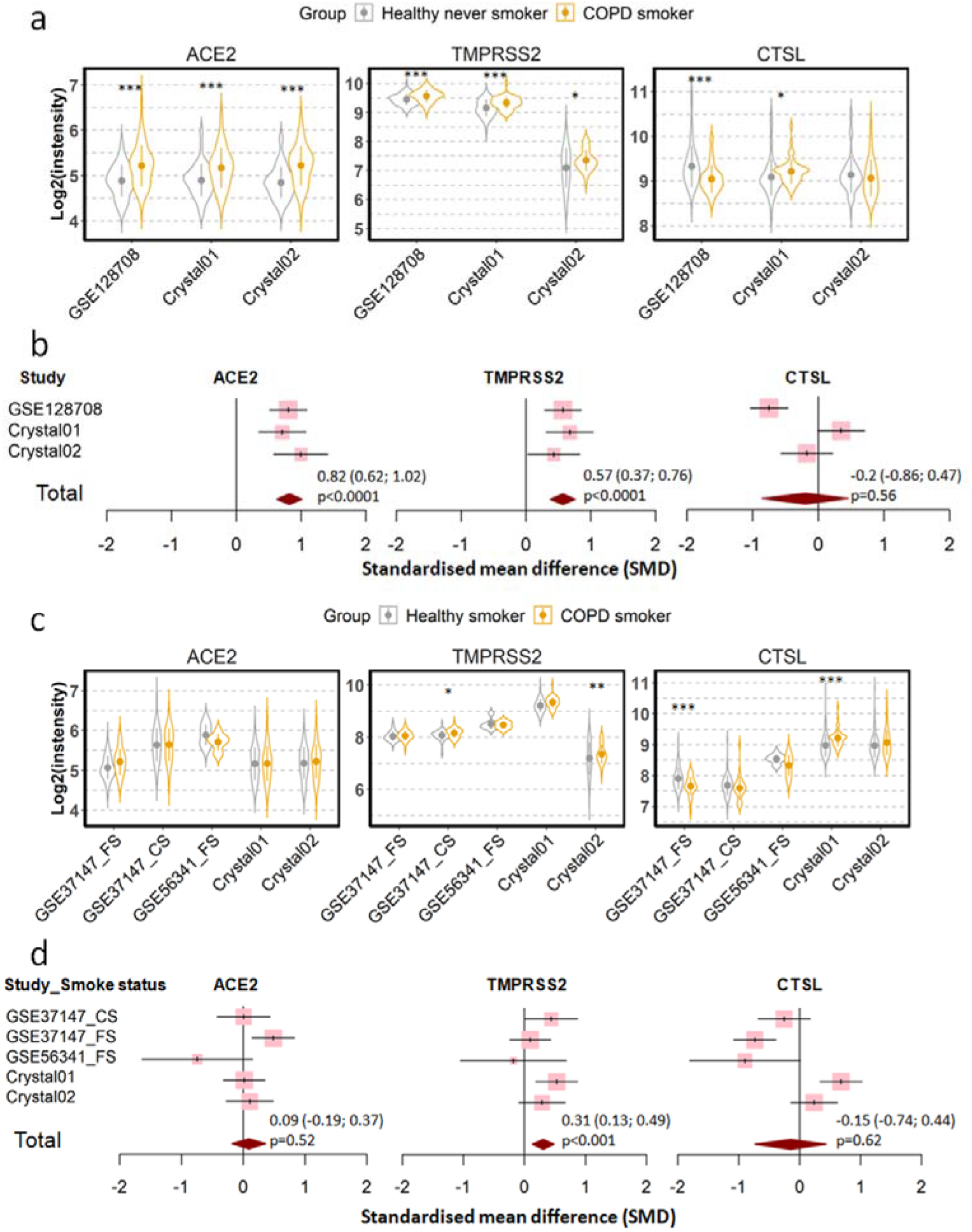
Expression of ACE2, TMPRSS2 and cathepsin L (CTSL) in airway epithelial cells of healthy subjects and COPD patients. **a)** Violin plot of expression levels of ACE2, TMPRSS2 and CTSL in 3 datasets which contain healthy never smokers and patients with COPD. Statistical difference was calculated by Student’s t test. *p<0.05, **p<0.01, ***p<0.001. **b)** Forest plot of 3 datasets examining expression of ACE2, TMPRSS2 and CTSL in healthy never smokers and patients with COPD. The SMD, 95% CI and *P* values of meta-analysis are depicted. **c)** Violin plot of expression levels of ACE2, TMPRSS2 and CTSL in 5 datasets which contain healthy smokers and patients with COPD. **d)** Forest plot of 5 datasets examining expression of ACE2, TMPRSS2 and CTSL in healthy smokers and patients with COPD. CS, Current smokers, FS, Former smokers, SM, Smokers.

Since most the COPD patients were smokers, we next determined whether the effect of COPD on expression of our target genes is due to cigarette smoking by comparing COPD patients with healthy smokers ^15, 17–21, 25, 26, 28, 31, 34, 37^. As shown in Fig. 3c-d, expression levels of ACE2 and cathepsin L are similar between both groups. The only difference between the two groups could be assigned to the expression of TMPRSS2, where COPD patients showed moderately higher RNA levels than healthy smokers (SMD=0.31, P<0.001). According to these results, the observed difference in the expression of the three SARS-CoV-2 entry-related genes between COPD patients and healthy controls is mainly due to cigarette smoking but not to disease manifestation.

To substantiate these findings, we examined the relationship between the lung function of COPD patients and gene expression by data from a further study in which lung function parameters were included ^34^., Expression levels of ACE2, TMPRSS2 and cathepsin L were not correlated with the first second of forced expiration (FEV1) or FEV1/FVC (Forced vital capacity) suggesting that the impairment of lung function does not affect the expression SARS-CoV-2 entry related genes (Supplementary Fig. 5).

Cigarette smoking does not only promote COPD but represents also a main cause for lung cancer. Therefore, we evaluated next the expression the three genes in patients with lung cancer and in corresponding healthy controls. In total, 7 datasets were recruited for the comparison, including 1003 patients and 540 healthy controls ^29, 32, 38–42^. Of note, healthy control subjects and patients in each dataset were identical in smoking status, either never smokers, former smokers or current smokers. Meta-analysis showed that lung cancer is associated with a very mild effect on modulation of expression of ACE2 (SMD=−0.16, P=0.0038) and cathepsin L (SMD=−0.18, P=0.0011) (Supplementary Fig. 6).

To include a chronic respiratory disease which induction is independent of smoking we analyzed one dataset derived from patients with cystic fibrosis ^43^. Since ACE2 expression was not analyzed here, we could only compare the expression of TMPRSS2 and cathepsin L between patients and healthy controls. With five subjects in each group, no significant difference in the expression of the two genes was observed between patients and controls (Supplementary Fig. 7).

### Expression of ACE2 and cathepsin L is decreased in nasal epithelial cells in asthma and allergic rhinitis

In the next step we determined the expression profile of the three genes in two chronic allergic respiratory diseases, allergic asthma and allergic rhinitis. The comparison between patients with allergic rhinitis and healthy subjects involved 3 comparisons, including two with bronchial epithelial cells and one with nasal epithelial cells ^44, 45^. Meta-analysis revealed a moderate decreasing effect of asthma on ACE2 (SMD=−0.76, P=0.0015) and cathepsin L (SMD=−0.62, P=0.039) expression. Moreover, this effect was seen more prominent on nasal epithelial cells than on bronchial epithelial cells (Supplementary Fig. 8a,b). A decreased expression of cathepsin L was also observed in patients with asthma as compared to healthy controls (SMD=−0.52, P=0.0049) ^30, 44–46^. Comparable to samples derived from rhinitis patients, the effect was drastically stronger on nasal epithelial cells than bronchial epithelial cells (Supplementary Fig. 8c,d).

Since both asthma and rhinitis are allergic respiratory diseases, we combined the two diseases to examine their effect on nasal epithelial cells. As expected, the meta-analysis revealed that the two diseases are associated with a strong effect on decreasing expression of ACE2 (SMD=−1.25, P<0.001) and cathepsin L (SMD=−0.77, P=0.0034) (Fig. 4).

**Fig. 4.**
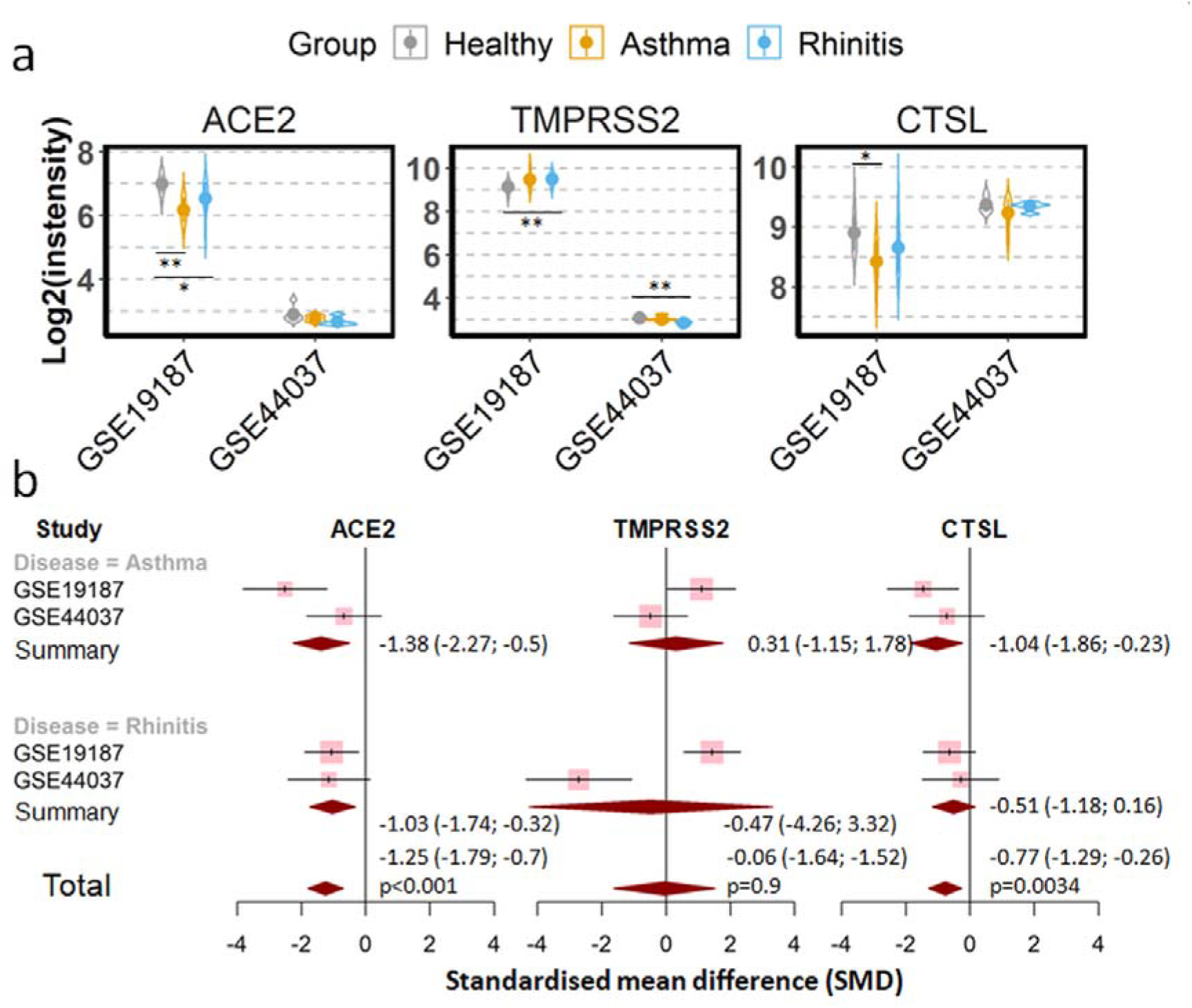
Decreased expression of ACE2 and cathepsin L (CTSL) in nasal epithelial cells of patients with allergic respiratory diseases. **a)** Violin plot of expression levels of ACE2, TMPRSS2 and CTSL in two datasets which contain healthy controls, patients with allergic rhinitis and patients with asthma. Statistical difference was calculated by Student’s t test. *p<0.05 and **p<0.01. **b)** Forest plot of datasets examining expression of ACE2, TMPRSS2 and CTSL in nasal epithelial cells of healthy controls and patients with allergic rhinitis or asthma. The x-axis indicates the standardized mean difference (SMD), while the y-axis shows GEO datasets and cell types. The SMD, 95% CI and *P* values of meta-analysis are depicted.

### Viral infection increases significantly the expression of ACE2, TMPRSS2 and cathepsin L

Finally, we examined the effect of viral infection, including experimental infection with rhinovirus in vivo and stimulation with rhinovirus in vitro ^47–49^, on the expression of the three SARS-CoV-2 entry related genes in airway epithelial cells.

As shown in Fig. 5, in vivo experimental infection with rhinovirus increased expression levels of ACE2, TMPRSS2 and cathepsin L 48 h after infection. In line with this result, in vitro stimulation of primary human airway epithelial cells isolated from healthy subjects or asthma patients with rhinovirus increased the expression of ACE2 and TMPRSS2 (Fig. 5).

**Fig. 5.**
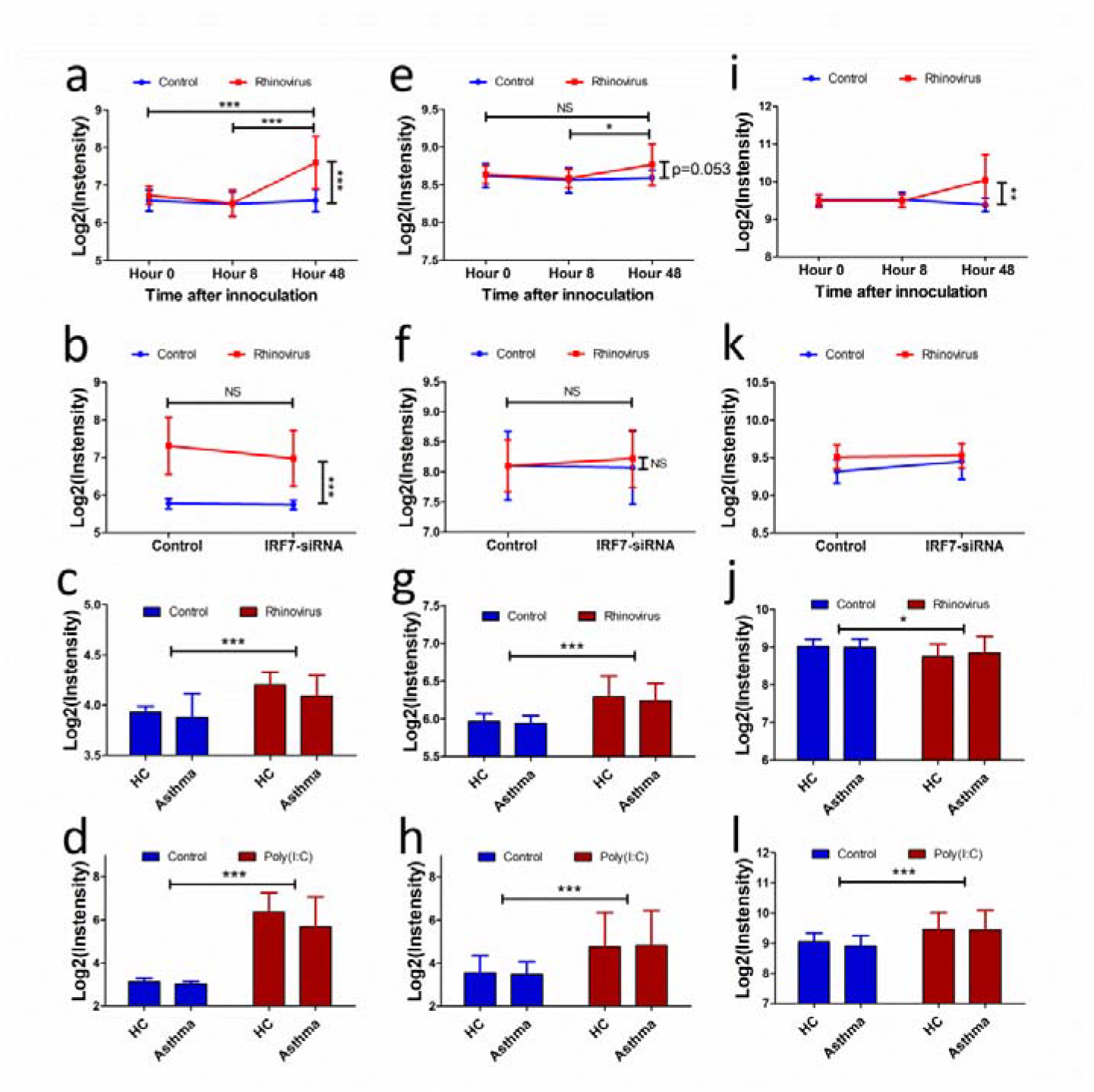
Expression of ACE2, TMPRSS2 and cathepsin L (CTSL) in airway epithelial cells after rhinoviral infection or TLR3 activation. Expression kinetics of of ACE2 (**a**), TMPRSS2 (**e**), and CTSL (**i**) in airway epithelial cells of healthy subjects experimentally infected with rhinovirus or saline-treated controls (data from GSE11348). Expression of ACE2 (**b**), TMPRSS2 (**f**), and CTSL (**j**) in airway epithelial cells isolated from healthy subjected and stimulated in vitro with or without rhinovirus in presence or absence of IRF-siRNA (data from GSE70190). Expression of ACE2 (**c**), TMPRSS2 (**g**), and CTSL (**k**) in airway epithelial cells isolated from healthy subjected or patients with asthma and stimulated in vitro with rhinovirus or saline control (data from GSE13396). Expression of ACE2 (**d**), TMPRSS2 (**h**), and CTSL (**l**) in airway epithelial cells isolated from healthy subjected or patients with asthma and stimulated in vitro with poly(I:C) or saline control (data from GSE13396). Statistical analysis was performed using two-way ANOVA, Tukey’s test for post hoc analysis was performed after two-way ANOVA analysis. *, p<0.05, **, p<0.01, ***, p<0.001. HC, healthy control, NS, not significant.

The stimulatory effect of viral infection on the expression of the SARS-CoV-2 entry associated genes was further confirmed in a dataset of human primary airway epithelial cells stimulated by poly(I:C) ^50^, a ligand of TLR3 which recognizes viral double-stranded RNA. After the stimulation, both primary epithelial cells from healthy subjects and patients with asthma expressed significantly higher levels of all three genes (Fig. 5).

Since it has been recently reported that ACE2 expression in airway epithelial cells is positively regulated by interferons ^51^, we speculated that this may reflect the mechanism underlying the effect of viral infection. To this end, we further examined the expression of IFNs in the datasets analyzed priviously. As shown in Supplementary Fig. 9, experimental viral infection, in vitro viral stimulation, and the poly(I:C) stimulation all resulted in an significantly increased expression of IFN-β supporting our notion.

## Discussion

In this study, we examined expression levels of ACE2, TMPRSS2 and cathepsin L in human airway epithelial cells derived from donors with different various physiological and pathological background. By comprehensive analysis, we generated a complete picture of expression profiling of the three SARS-CoV-2 entry related genes.

In our current study we can demonstrate that current smokers show a highly significant increase in ACE2 expression as compared to never smokers, which is in line with previous findings ^14^. Moreover, our study extends the analysis to two other SARS-CoV-2 entry-related genes, TMPRSS2 and cathepsin L. Unexpectedly, smoking exerts antagonistic effects on the expression of TMPRSS2 and cathepsin L in airway epithelial cells by increasing the expression of the former but decreasing the latter. In several recent meta-analysis it was reported that smoking is associated with an approximately 2 fold increase in the risk of developing severe COVID-19 ^12, 52, 53^, and it has been hypothesized that this increase may be caused by an elevated expression of ACE2 in smoking subjects^14^. However, our data implicate that the disease-promoting effect of smoking cannot be explained by a simple up-regulation of genes identified to be relevant for the virus entry. First of all, the difference between never-smokers and current smokers appears with a 20% elevation in ACE2 mRNA expression in the latter group as rather low and is unclear, whether this minor disparity has any pathophysiological consequence. Moreover, smoking increases the expression of TMPRSS2 but decreases the expression of cathepsin L. Accordingly, it is conceivable that smoking might enhance the entry of SARS-CoV-2 via membrane fusion but decrease the entry via endocytosis. Since it remains unknown whether the viral entry via the two ways make difference for SARS-CoV-2 infection ^7, 54^, the mechanisms of smoking on SARS-CoV-2 infection remain to be elucidated.

Irrespective of the unclear pathomechanism of smoking on SARS-CoV-2 infection, our study uncovered an unexpected difference in the expression of the three genes between current smokers and former smokers, but not between former smokers and never smokers. This result indicates an enormous acute effect of smoking on the expression of SARS-CoV-2 entry-associated genes. In contrast to many chronic and irreversible changes in the lung, smoking cessation could be effective to reduce the potential risk emanating from the alternation of the viral entry associated genes.

According to our findings, COPD, a chronic respiratory disease mainly caused by cigarette smoking, is also associated with an increased expression of ACE2 in human airway epthelial cells. However, the effect seems largely related to smoking rather than the disease itself since the differences of the gene expression become visible only between COPD patients and never-smoker healthy controls, but do not appear between COPD patients and healthy smoker controls, ii) the variations in the expression of the three genes between COPD patients and never smoker controls follow the same pattern as that between healthy smokers and never smokers, and iii) lung function is not correlated with the expression of the three genes.

Similar to COPD, patients with lung cancer also do not show high levels of the three SARS-CoV-2 related genes than healthy smoker controls. In addition, those genes are also not upregulated airway epithelial cells of three other respiratory diseases, namely asthma, allergic rhinitis and cystic fibrosis. Taken together, these results do not support the hypothesis that the high susceptibility to severe COVID-19 in patients with chronic respiratory diseases could be explained by a higher expression of SARS-CoV-2 entry genes in those patients as compared to controls.

Notably, the expression of ACE2 and cathepsin L is significantly and considerably decreased in nasal epithelial cells of patients with allergic respiratory disease, including both allergic asthma and allergic rhinitis, compared with healthy controls. This observation is in line with findings in recent two studies in which ACE2 expression in airway epithelial cells in patient with asthma and healthy subjects were reported ^55, 56^. Taken together, this suggests that type 2 inflammatory processes might modulate the expression of SARS-CoV-2 entry related genes. Accordingly, it is conceivable that patients with allergic respiratory diseases might be less susceptible to SARS-CoV-2 infection. However, this speculation needs to be verified by investigating the prevalence of COVID-19 in those populations. Also, further exploration of the role of type 2 immune responses in SARS-CoV-2 infection could help to control the pandemic.

The most striking effect on the expression of ACE2, TMPRSS2 and cathepsin L was observed in rhinovirus infection. Both experimental infection with rhinovirus in vivo and stimulation with virus in vitro led to a dramatic upregulation of the ACE2 expression. Moreover, poly(I:C) which mimics the viral stimulation also shows a similar effect. Recently, Ziegler et al. demonstrated that ACE2 is an interferon stimulated gene in human airway epithelial cells ^51^. The current study also demonstrates that the expression of IFN-ß in airway epithelial cells is highly upregulated after rhinovirus infection or poly(I:C) stimulation, supporting that ACE2, as well as TMPRSS2 and cathepsin L are interferon stimulated genes. Since rhinoviruses are major pathogens of common colds ^57^, it will be interesting to determine the effect of common cold on the susceptibility and severity of COVID-19.

In conclusion, this study provides an overview of the expression profile of SARS-CoV-2 entry associated genes in human airway epithelial cells under various physiological and pathological conditions. The results from this study implicate that respiratory viral infection, cigarette smoking and allergic respiratory diseases might affect the entry of the SARS-CoV-2 and consequent development of COVID-19.

## Methods

### Data retrieval

An exhaustive search of the Gene Expression Omnibus (GEO) database (https://www.ncbi.nlm.nih.gov/geo/) was performed to identify eligible data on 1st June 2020. The procedure of identification of eligible data is shown in Figure 1. First of all, the key word “Airway epithelial cells” was used for the search without any limitations. In a second step, non-human datasets, non-series datasets and non-gene expression array datasets were filtered out. Finally, datasets from step two were reviewed carefully, and the following datasets were excluded: (a) datasets of cell line(s), (b) redundant dataset, (c) datasets containing only one group and (d) datasets containing less than 5 samples per group.

### Data correction, normalization

For affymetrix microarrays, CEL files were uploaded into RStudio (Version 1.3.959, based on R version 4.0.1) using package “affy” ^58^. Subsequently, background correction and normalization were applied to the raw data using “Robust Multichip Average (RMA)” method 59. For agilent microarray data, raw expression profiling files were uploaded into RStudio using package “limma” ^60^. By using the “limma” package, background correction and normalization were performed using “normexp” and “quantile” methods, respectively. Background corrected, normalized and log2 transformed signal intensities of ACE2, TMPRSS2 and cathepsin L were outputted and used for further analysis.

### Meta analysis

All meta-analysis were performed using “meta” package in RStudio. Standardised mean difference (SMD) was utilized to assess the effect size of a factor on the expression of targeted genes, and 95% confidence intervals (CIs) of SMD was calculated ^61^. According to the guideline proposed by Cohen^62^, the magnitude of the SMD is interpreted as below: small, SMD = 0.2; medium, SMD = 0.5; and large, SMD = 0.8. Fixed or random effect model was applied to pool the effect size depending on the heterogeneity across the datasets determined by inconsistency (I^2^) statistics and Cochrane’s Q test. random effect model was applied when there was significant heterogeneity among datasets (I^2^ value > 50% or *P* value of Q test < 0.05), otherwise fixed effect model was utilized ^63, 64^.

### Statistics

All statistical analyses were conducted using RStudio. Statistical significance between two groups was calculated using paired or unpaired Student’s t test depending on the samples. For the study containing two categorical factors, we utilized two-way ANOVA to determine the statistical difference followed by Tukey’s test for post hoc analysis. Multiple linear regression model was generated to evaluate the correlation of lung function index of COPD patients and healthy controls with gene expression of ACE2. A *P* value<0.05 was considered as statistical significance.

## Author Contribution

Study design: X.Y., Literature search and data analysis: J.Y., Data interpretation: B.K. F.P. X.Y., Writing: B.K. F.P. X.Y.

## Reference List

1. Huang,C. et al. Clinical features of patients infected with 2019 novel coronavirus in Wuhan, China. Lancet 395, 497–506 (2020).

2. Zhu,N. et al. A Novel Coronavirus from Patients with Pneumonia in China, 2019. N. Engl. J. Med.(2020).

3. WHO characterizes COVID-19 as a pandemic. 2020. Ref Type: Internet Communication

4. COVID-19 Coronavirus Pandemic. 2020. Ref Type: Internet Communication

5. Lu,R. et al. Genomic characterisation and epidemiology of 2019 novel coronavirus: implications for virus origins and receptor binding. Lancet(2020).

6. Zhou,P. et al. A pneumonia outbreak associated with a new coronavirus of probable bat origin. Nature(2020).

7. Heurich,A. et al. TMPRSS2 and ADAM17 cleave ACE2 differentially and only proteolysis by TMPRSS2 augments entry driven by the severe acute respiratory syndrome coronavirus spike protein. J. Virol. 88, 1293–1307 (2014).

8. Hoffmann,M. et al. SARS-CoV-2 Cell Entry Depends on ACE2 and TMPRSS2 and Is Blocked by a Clinically Proven Protease Inhibitor. Cell 181, 271–280 (2020).

9. Ou,X. et al. Characterization of spike glycoprotein of SARS-CoV-2 on virus entry and its immune cross-reactivity with SARS-CoV. Nat. Commun. 11, 1620 (2020).

10. Guan,W.J. et al. Clinical Characteristics of Coronavirus Disease 2019 in China. N. Engl. J. Med. 382, 1708–1720 (2020).

11. Lippi,G. & Henry,B.M. Chronic obstructive pulmonary disease is associated with severe coronavirus disease 2019 (COVID-19). Respir. Med. 167, 105941 (2020).

12. Zhao,Q. et al. The impact of COPD and smoking history on the severity of COVID-19: A systemic review and meta-analysis. J. Med. Virol.(2020).

13. Zhang,M., Gao,Y., Zhao,W., Yu,G., & Jin,F. ACE-2/ANG 1-7 ameliorates ER stress-induced apoptosis in seawater aspiration-induced acute lung injury. Am. J. Physiol Lung Cell Mol. Physiol 315, L1015–L1027 (2018).

14. Leung,J.M. et al. ACE-2 expression in the small airway epithelia of smokers and COPD patients: implications for COVID-19. Eur. Respir. J. 55, (2020).

15. Ammous,Z. et al. Variability in small airway epithelial gene expression among normal smokers. Chest 133, 1344–1353 (2008).

16. Beane,J. et al. Reversible and permanent effects of tobacco smoke exposure on airway epithelial gene expression. Genome Biol. 8, R201 (2007).

17. Butler,M.W. et al. Modulation of cystatin A expression in human airway epithelium related to genotype, smoking, COPD, and lung cancer. Cancer Res. 71, 2572–2581 (2011).

18. Carolan,B.J. et al. Up-regulation of expression of the ubiquitin carboxyl-terminal hydrolase L1 gene in human airway epithelium of cigarette smokers. Cancer Res. 66, 10729–10740 (2006).

19. Carolan,B.J., Harvey,B.G., De,B.P., Vanni,H., & Crystal,R.G. Decreased expression of intelectin 1 in the human airway epithelium of smokers compared to nonsmokers. J. Immunol. 181, 5760–5767 (2008).

20. Hessel,J. et al. Intraflagellar transport gene expression associated with short cilia in smoking and COPD. PLoS. One. 9, e85453 (2014).

21. Raman,T. et al. Quality control in microarray assessment of gene expression in human airway epithelium. BMC. Genomics 10, 493 (2009).

22. Schembri,F. et al. MicroRNAs as modulators of smoking-induced gene expression changes in human airway epithelium. Proc. Natl. Acad. Sci. U. S. A 106, 2319–2324 (2009).

23. Spira,A. et al. Effects of cigarette smoke on the human airway epithelial cell transcriptome. Proc. Natl. Acad. Sci. U. S. A 101, 10143–10148 (2004).

24. Sridhar,S. et al. Smoking-induced gene expression changes in the bronchial airway are reflected in nasal and buccal epithelium. BMC. Genomics 9, 259 (2008).

25. Strulovici-Barel,Y. et al. Threshold of biologic responses of the small airway epithelium to low levels of tobacco smoke. Am. J. Respir. Crit Care Med. 182, 1524–1532 (2010).

26. Tilley,A.E. et al. Biologic phenotyping of the human small airway epithelial response to cigarette smoking. PLoS. One. 6, e22798 (2011).

27. Walters,M.S. et al. Waterpipe smoking induces epigenetic changes in the small airway epithelium. PLoS. One. 12, e0171112 (2017).

28. Wang,R. et al. Airway epithelial expression of TLR5 is downregulated in healthy smokers and smokers with chronic obstructive pulmonary disease. J. Immunol. 189, 2217–2225 (2012).

29. Wang,X. et al. Genetic variation and antioxidant response gene expression in the bronchial airway epithelium of smokers at risk for lung cancer. PLoS. One. 5, e11934 (2010).

30. Woodruff,P.G. et al. Genome-wide profiling identifies epithelial cell genes associated with asthma and with treatment response to corticosteroids. Proc. Natl. Acad. Sci. U. S. A 104, 15858–15863 (2007).

31. Zhou,H. et al. POU2AF1 Functions in the Human Airway Epithelium To Regulate Expression of Host Defense Genes. J. Immunol. 196, 3159–3167 (2016).

32. Shared Gene Expression Alterations in Nasal and Bronchial Epithelium for Lung Cancer Detection. J. Natl. Cancer Inst. 109, (2017).

33. Corbett,S.E. et al. Gene Expression Alterations in the Bronchial Epithelium of e-Cigarette Users. Chest 156, 764–773 (2019).

34. Travis,W.D. et al. The 2015 World Health Organization Classification of Lung Tumors: Impact of Genetic, Clinical and Radiologic Advances Since the 2004 Classification. J. Thorac. Oncol. 10, 1243–1260 (2015).

35. Billatos,E. et al. Impact of acute exposure to cigarette smoke on airway gene expression. Physiol Genomics 50, 705–713 (2018).

36. Zhang,H. et al. Expression of the SARS-CoV-2 ACE2 Receptor in the Human Airway Epithelium. Am. J. Respir. Crit Care Med.(2020).

37. Vucic,E.A. et al. DNA methylation is globally disrupted and associated with expression changes in chronic obstructive pulmonary disease small airways. Am. J. Respir. Cell Mol. Biol. 50, 912–922 (2014).

38. Beane,J. et al. Characterizing the impact of smoking and lung cancer on the airway transcriptome using RNA-Seq. Cancer Prev. Res. (Phila) 4, 803–817 (2011).

39. Silvestri,G.A. et al. A Bronchial Genomic Classifier for the Diagnostic Evaluation of Lung Cancer. N. Engl. J. Med. 373, 243–251 (2015).

40. Spira,A. et al. Airway epithelial gene expression in the diagnostic evaluation of smokers with suspect lung cancer. Nat. Med. 13, 361–366 (2007).

41. Tong,R. et al. Decreased Interferon Alpha/Beta Signature Associated with Human Lung Tumorigenesis. J. Interferon Cytokine Res. 35, 963–968 (2015).

42. Tsay,J.C. et al. Molecular characterization of the peripheral airway field of cancerization in lung adenocarcinoma. PLoS. One. 10, e0118132 (2015).

43. Clarke,L.A., Sousa,L., Barreto,C., & Amaral,M.D. Changes in transcriptome of native nasal epithelium expressing F508del-CFTR and intersecting data from comparable studies. Respir. Res. 14, 38 (2013).

44. Giovannini-Chami,L. et al. Distinct epithelial gene expression phenotypes in childhood respiratory allergy. Eur. Respir. J. 39, 1197–1205 (2012).

45. Wagener,A.H. et al. The impact of allergic rhinitis and asthma on human nasal and bronchial epithelial gene expression. PLoS. One. 8, e80257 (2013).

46. Kicic,A. et al. Decreased fibronectin production significantly contributes to dysregulated repair of asthmatic epithelium. Am. J. Respir. Crit Care Med. 181, 889–898 (2010).

47. Bochkov,Y.A. et al. Rhinovirus-induced modulation of gene expression in bronchial epithelial cells from subjects with asthma. Mucosal. Immunol. 3, 69–80 (2010).

48. Bosco,A., Wiehler,S., & Proud,D. Interferon regulatory factor 7 regulates airway epithelial cell responses to human rhinovirus infection. BMC. Genomics 17, 76 (2016).

49. Proud,D. et al. Gene expression profiles during in vivo human rhinovirus infection: insights into the host response. Am. J. Respir. Crit Care Med. 178, 962–968 (2008).

50. Wagener,A.H. et al. dsRNA-induced changes in gene expression profiles of primary nasal and bronchial epithelial cells from patients with asthma, rhinitis and controls. Respir. Res. 15, 9 (2014).

51. Ziegler,C.G.K. et al. SARS-CoV-2 Receptor ACE2 Is an Interferon-Stimulated Gene in Human Airway Epithelial Cells and Is Detected in Specific Cell Subsets across Tissues. Cell 181, 1016–1035 (2020).

52. Guo,F.R. Active smoking is associated with severity of coronavirus disease 2019 (COVID-19): An update of a meta-analysis. Tob. Induc. Dis. 18, 37 (2020).

53. Zheng,Z. et al. Risk factors of critical & mortal COVID-19 cases: A systematic literature review and meta-analysis. J. Infect.(2020).

54. Pillay,T.S. Gene of the month: the 2019-nCoV/SARS-CoV-2 novel coronavirus spike protein. J. Clin. Pathol. 73, 366–369 (2020).

55. Camiolo,M.J., Gauthier,M., Kaminski,N., Ray,A., & Wenzel,S.E. Expression of SARS-CoV-2 Receptor ACE2 and Coincident Host Response Signature Varies by Asthma Inflammatory Phenotype. J. Allergy Clin. Immunol.(2020).

56. Jackson,D.J. et al. Association of respiratory allergy, asthma, and expression of the SARS-CoV-2 receptor ACE2. J. Allergy Clin. Immunol.(2020).

57. Bashir,H. et al. Association of rhinovirus species with common cold and asthma symptoms and bacterial pathogens. J. Allergy Clin. Immunol. 141, 822–824 (2018).

58. Gautier,L., Cope,L., Bolstad,B.M., & Irizarry,R.A. affy--analysis of Affymetrix GeneChip data at the probe level. Bioinformatics. 20, 307–315 (2004).

59. Irizarry,R.A. et al. Exploration, normalization, and summaries of high density oligonucleotide array probe level data. Biostatistics. 4, 249–264 (2003).

60. Ritchie,M.E. et al. limma powers differential expression analyses for RNA-sequencing and microarray studies. Nucleic Acids Res. 43, e47 (2015).

61. Faraone,S.V. Interpreting estimates of treatment effects: implications for managed care. P. T. 33, 700–711 (2008).

62. Cohen J. Statistical Power Analysis for the Behavioral Sciences.1988).

63. He,R.Q. et al. Downregulated miR-23b-3p expression acts as a predictor of hepatocellular carcinoma progression: A study based on public data and RT-qPCR verification. Int. J. Mol. Med. 41, 2813–2831 (2018).

64. Higgins,J.P. & Thompson,S.G. Quantifying heterogeneity in a meta-analysis. Stat. Med. 21, 1539–1558 (2002).

