## Supplementary tables and figures for "Expression of ACE2, TMPRSS2 and CTSL in human airway epithelial cells under physiological and pathological conditions: Implications for SARS-CoV-2 infection"

### Supplementary information

#### Supplementary table 1: List of datasets recruited in this study

| Datasets | Platform | Ref. (PMID) | Studying focus | Submitted time (year) | Cell type | Sample sizes |
|---|---|---|---|---|---|---|
| Crystal et al. Group 01 (GSE4498, GSE5058, GSE5059, GSE7832, GSE8545, GSE10006, GSE10135, GSE11784, GSE11906, GSE11952, GSE13931, GSE13933, GSE17905, GSE18385, GSE19407, GSE19667, GSE20250, GSE20257) | GPL570 [HG-U133_Plus_2] | 18339782, 17108109, 18832735, 21829517, 19852842, 20693378 | Smoking and COPD | 2006-2010 | AEC | 49 COPD, 124 NS ,167 CS |
| Crystal et al. Group 02 (GSE22047, GSE24337, GSE30063, GSE34450, GSE43939, GSE53537, GSE63127, GSE64614, GSE76324, GSE77658) | GPL570 [HG-U133_Plus_2] | 21325429, 22855713, 24465567, 26927796 | Smoking and COPD | 2010-2016 | AEC | 36 COPD, 98 NS, 129 CS |
| GSE994 | GPL96 [HG-U133A] | 15210990 | Smoking | 2004 | BEC | 34 CS, 18 FS, 23 NS |
| GSE4115 | GPL96 [HG-U133A] | 17334370 | Lung cancer | 2006 | BEC | 102 LC, 90 NC |
| GSE4302 | GPL570 [HG-U133_Plus_2] | 17898169 | Smoking and athma | 2006 | BEC | 42 baseline asthma, 13 placebo treated asthma, 19 Flovent treated asthma, 16 smoker, 28 HC |
| GSE7895 | GPL96 [HG-U133A] | 17894889 | Smoking | 2007 | BEC | 52 CS, 31 FS, 21 NS |
| GSE8987 | GPL571 [HG-U133A_2] | 18513428 | Smoking | 2007 | NEC | 7 CS, 8 NS |
| GSE11348 | GPL570 [HG-U133_Plus_2] | 18658112 | Human rhinovirus infection | 2008 | NEC | 48 Control 45 HRV |
| GSE13396 | GPL570 [HG-U133_Plus_2] | 19710636 | Human rhinovirus infection | 2008 | BEC | 11 Control 11 HRV |
| GSE14633 | GPL5175 [HuEx-1_0-st] | 19168627 | Smoking | 2009 | BEC | 11 CS, 11 NS |
| GSE18965 | GPL96 [HG-U133A] | 20110557 | Asthma | 2009 | BEC | 9 Asthma, 7 HC |
| GSE19027 | GPL96 [HG-U133A] | 20689807 | Smoking and lung cancer | 2009 | BEC | CA: 9 CS, 12 FS; HC: 20 CS 9 FS 9 NS |
| GSE19187 | GPL6244 [HuGene-1_0-st] | 22005912 | Allergic rhinitis and asthma | 2009 | NEC | 7 Controled asthma,, 6 uncontroled asthma, 11 HC, 14 Rhinitis |
| GSE28835 | GPL13447 [HG-U133A_2] | 21636547 | Lung cancer | 2011 | LAEC | 8 LC ,5 NC |
| GSE37147 | GPL13243 [HuGene10stv1_Hs_ENSG] | 23471465 | COPD | 2012 | BEC | COPD: 30 CS, 57 FS; HC: 69 CS, 82 FS |
| GSE40445 | GPL10097 [HsAirwaya520108F] | 23537407 | Cystic fibrosis | 2012 | NEC | 5 CF, 5 HC |
| GSE44037 | GPL13158 [HT_HG-U133_Plus_PM] | 24282527 | Allergic rhinitis and asthma | 2013 | BEC | 12 Asthma ,12 HC, 10 Rhinitis |
| GSE51392 | GPL13158 [HT_HG-U133_Plus_PM] | 24475887 | Poly(I:C) stimulation | 2013 | BEC | 34 Control 34 Poly(I:C) |
| GSE54495 | GPL570 [HG-U133_Plus_2] | 25705890 | Lung cancer | 2014 | Peripheral AEC | 17 LC, 13 Smoker |

| | | | | | | |
|---|---|---|---|---|---|---|
| GSE56341 | GPL6244 [HuGene-1_0-st] | 24298892 | COPD | 2014 | Small AEC | 8 COPD, 14 FS |
| GSE66499 | GPL6244 [HuGene-1_0-st] | 25981554 | Lung cancer | 2015 | BEC | 490 LC ,190 HC |
| GSE67061 | GPL17077 [Agilent-039494] | 26308599 | Lung cancer | 2015 | AEC | 56 LC ,17 HC |
| GSE70190 | GPL20609 [PrimeView] | 26810609 | Human rhinovirus infection | 2015 | BEC | 10 Control, 10 rhinovirus infection |
| GSE80796 | GPL6244 [HuGene-1_0-st] | 28376173 | Smoking and lung cancer | 2016 | NEC | CA: 113 CS, 196 FS; HC: 73 CS, 123 FS |
| GSE84101 | GPL570 [HG-U133_Plus_2] | 28273093 | Smoking | 2016 | Small AEC | 7 SM ,7 NS |
| GSE97010 | GPL17244 [HuGene-1_0-st] | 29932825 | Smoking | 2017 | BEC | 63 baseline ,63 post ASE |
| GSE112073 | GPL17556 [HuGene-1_0-st] | 31233743 | Smoking | 2018 | BEC | 9 CS, 21 FS |
| GSE128708 | GPL570 [HG-U133_Plus_2] | 32432483 | COPD | 2019 | Small AEC | 124 COPD smoker, 84 NS |

AEC, Airway epithelial cells; BEC, Bronchial and nasal epithelial cells; LAEC, Large airway epithelial cells; NEC, Nasal epithelial cells; NS, Never smoker; SM, Smoker; CS, Current smoker; FS, Former smoker; LC, Lung cancer; HC, Healthy control; COPD,. Chronic obstractive pulmonary disease; ASE, Acute smoking exposure.

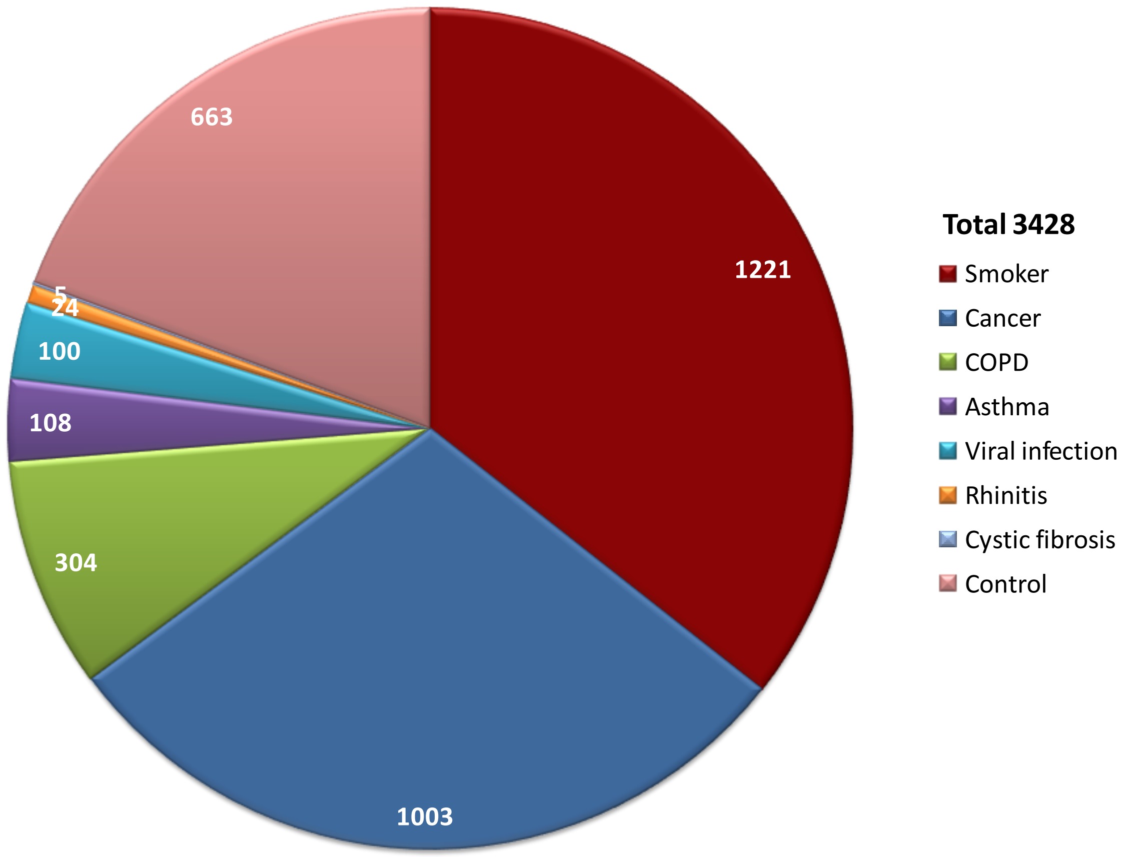


**Supplementary fig.1: Pie chart showing the composition of samples.** Control group consists of healthy never smokers, healthy subjects without information of smoking status and control samples in *in vitro* studies, while the smoker group contains healthy smokers, including both current smokers and former smokers.

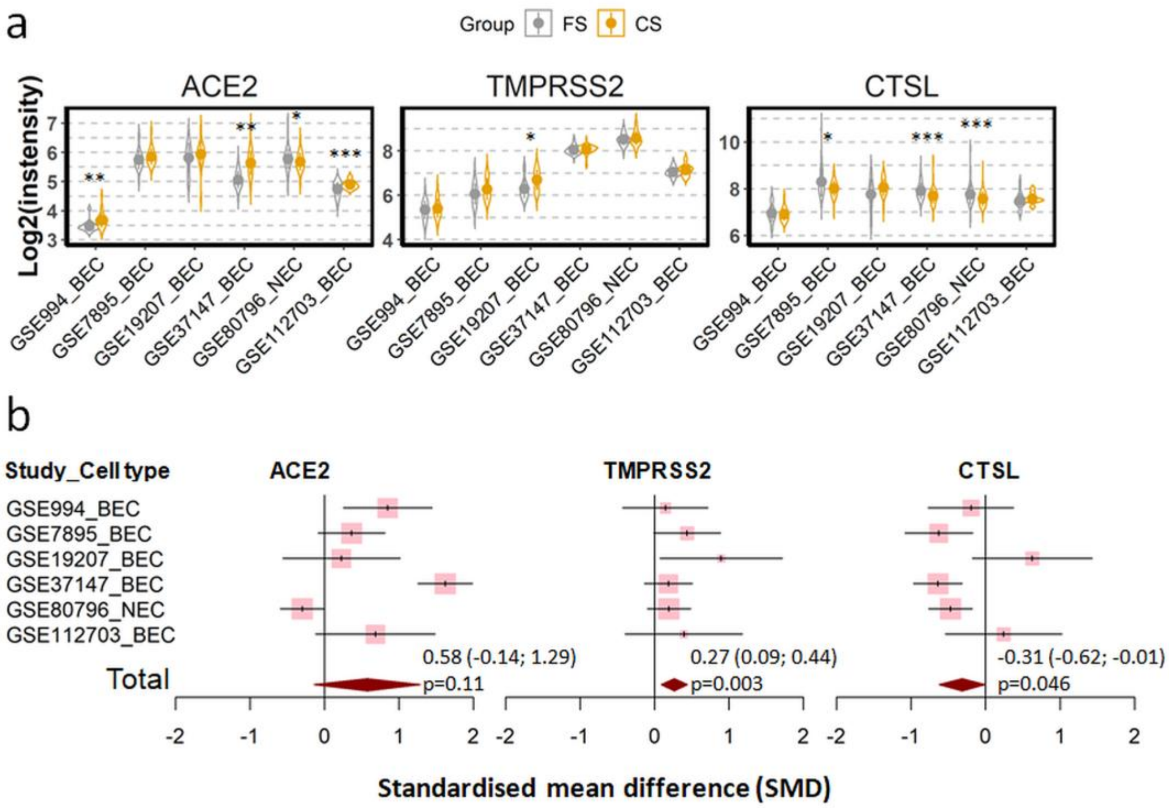


**Supplementary Fig. 2. Expression of ACE2, TMRPSS2 and cathepsin L (CTSL) in airway epithelial cells of healthy current smokers (CS) and former smokers (FS). a)** Violin plot of expression levels of ACE2, TMPRSS2 and CTSL in 6 datasets which contain current smokers and former smoker. The mean and standard deviation (SD) of each group were presented as dot and line, respectively. BEC, bronchial epithelial cells; NEC, nasal epithelial cells. Statistical differences were calculated by Student's t test. *p<0.05, **p<0.01, ***p<0.001. **b)** Forest plot of 6 datasets examining expression of ACE2, TMPRSS2 and CTSL in current smokers and former smokers. The x-axis indicates the standardized mean difference (SMD), while the y-axis shows GEO datasets and cell types. Each square in the plots represents the SMD in corresponding datasets and the 95% confidence interval (CI) is showed by the error bar. The size of each square represents the weight of the individual dataset in the meta-analysis. The diamonds in the bottom represent the SMD of the meta-analysis. The SMD, 95% CI and *P* values of meta-analysis are depicted.

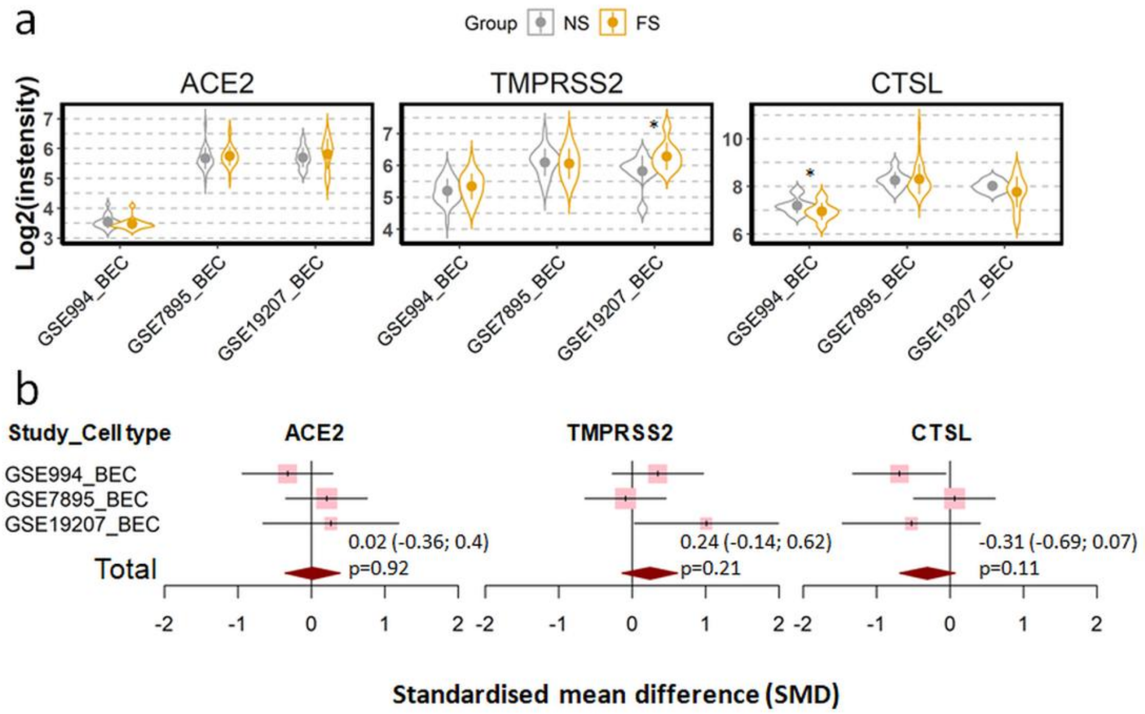


**Supplementary Fig. 3. Expression of ACE2, TMRPSS2 and cathepsin L (CTSL) in the airway epithelial cells of healthy never smokers (NS) and former smokers (FS) . a)** Violin plot of expression levels of ACE2, TMPRSS2 and CTSL in 3 datasets which contain current smokers and former smoker. Statistical difference was calculated by Student's t test. *p<0.05. **b)** Forest plot of 3 datasets examining expression of ACE2, TMPRSS2 and CTSL in never smokers and former smokers. The x-axis indicates the standardized mean difference (SMD), while the y-axis shows GEO datasets and cell types. The SMD, 95% CI and *P* values of meta-analysis are depicted.

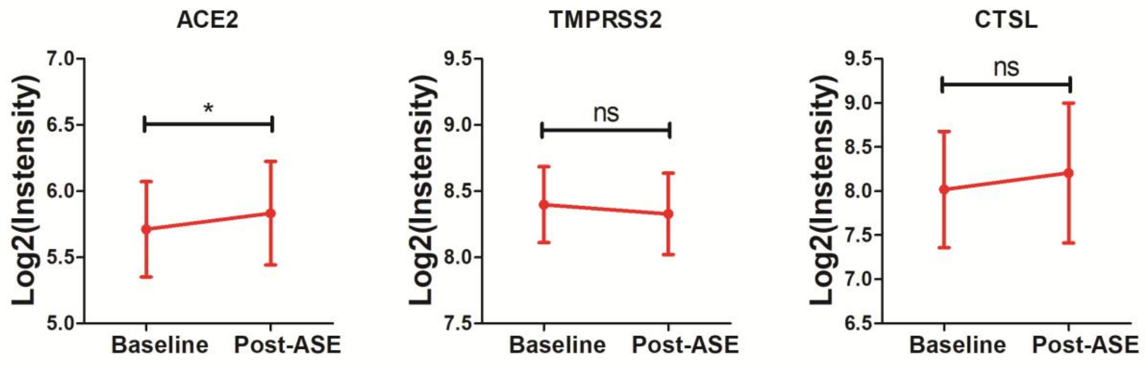


**Supplementary Fig. 4. Expression of ACE2, TMPRSS2 and cathepsin L (CTSL) in airway epithelial cells before and after acute smoke exposure (ASE) in healthy smokers in dataset GSE97010.** The post-ASE group stands for 63 smokers who were asked to refrain from cigarette smoking for at least 2 days and then subjected to ASE, while the 63 subjects obtained from smoking and underwent bronchoscopy at a separate time at least 6 wks from the post-smoking bronchoscopy to serve as an unexposed baseline group. Statistical difference was analyzed by paired Student t test. * $p<0.05$, ns. not significant.

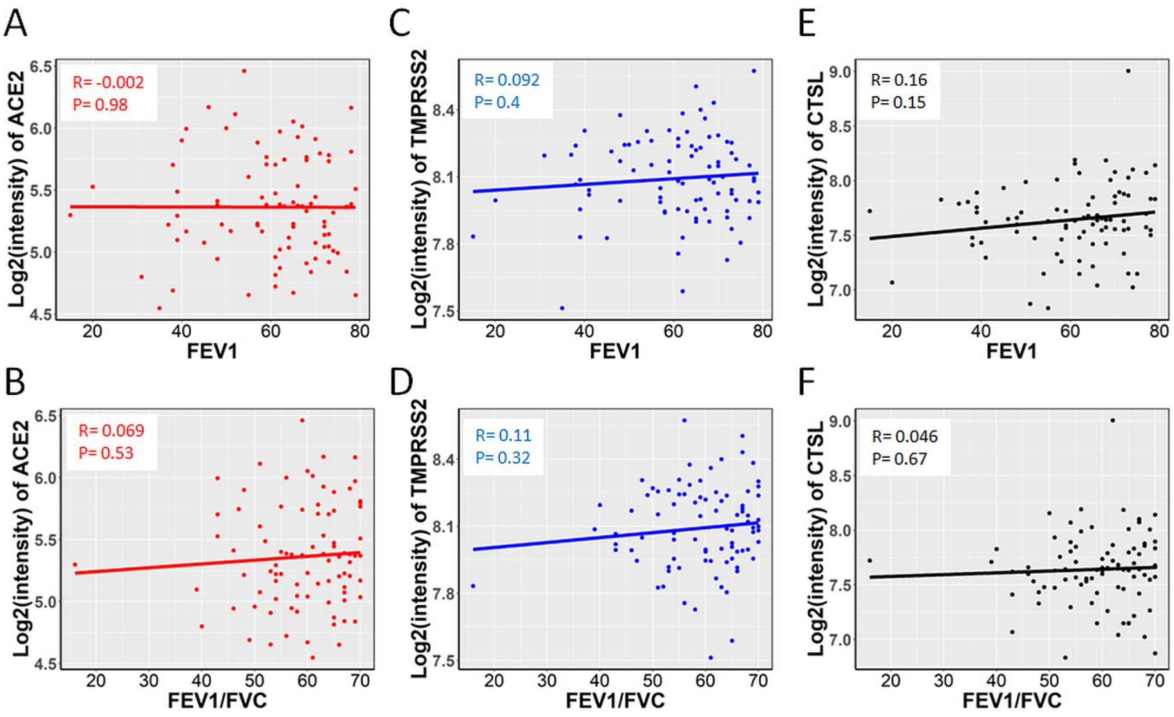


**Supplementary Fig. 5. Correlation between SARS-CoV-2 entry related genes with lung function indexes in COPD patients.** Linear regression of FEV1 (A,C,E) and FEV1/FVC (B,D,F) with expression of ACE2 (A,B), TMPRSS2 (C,D) and cathepsin L (CTSL) (E,F). Data are from dataset GSE37147. FEV1, the first second of forced expiration to the full; FVC, forced vital capacity; FEV1/FVC, ratio of FEV1 and FVC. *P* values and correlation coefficient (R) calculated by pearson method are depicted.

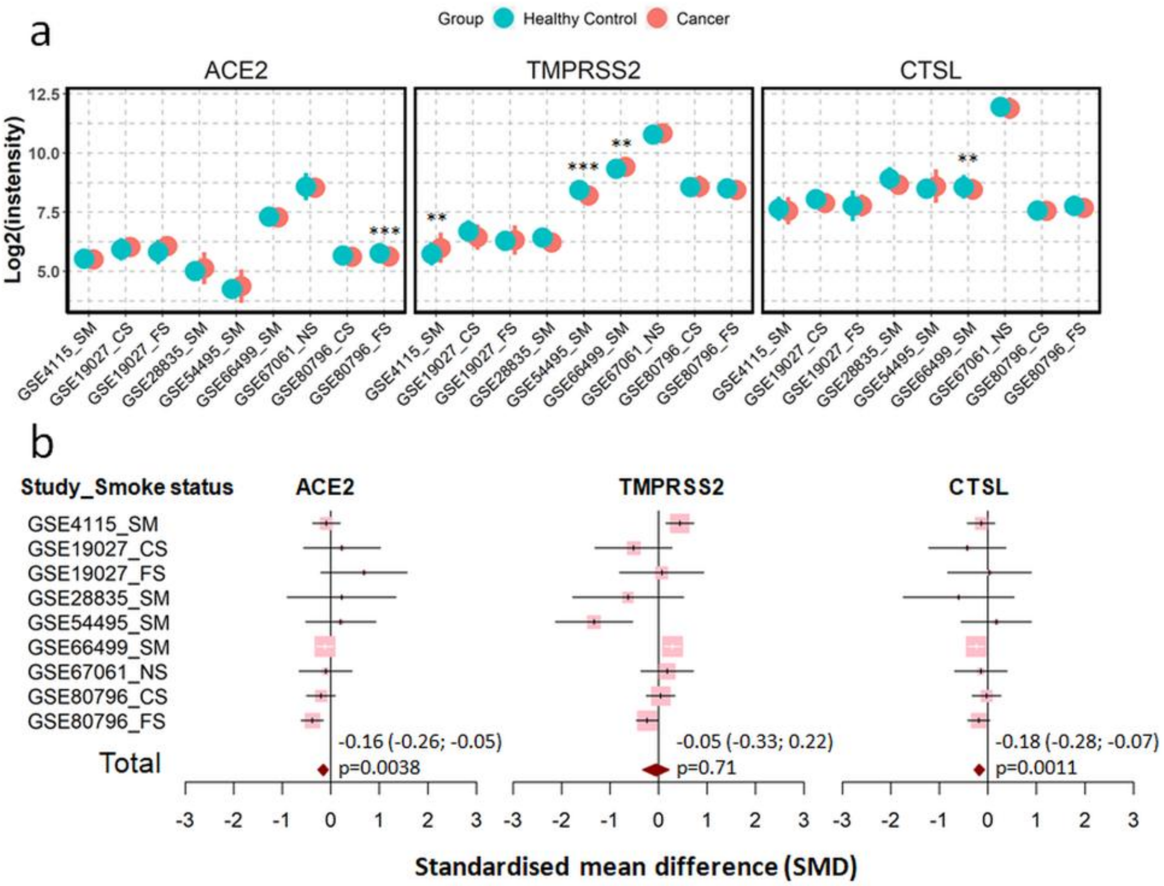


**Supplementary Fig. 6. Expression of ACE2, TMRPSS2 and cathepsin L (CTSL) in airway epithelial cells of healthy controls and patients with lung cancer. a)** Plot of expression levels of ACE2, TMPRSS2 and CTSL in 9 datasets which contain healthy control and patients with lung cancers. Statistical difference was calculated by Student's t test. *p<0.05, **p<0.01, ***p<0.001. **b)** Forest plot of 9 datasets examining expression of ACE2, TMPRSS2 and CTSL in healthy controls and patients with lung cancer. The x-axis indicates the standardized mean difference (SMD), while the y-axis shows GEO datasets and cell types. The SMD, 95% CI and p values of meta-analysis are depicted. BEC, bronchial epithelial cells; NEC, nasal epithelial cells; CS, Current smokers; FS, Former smokers; SM, Smokers.

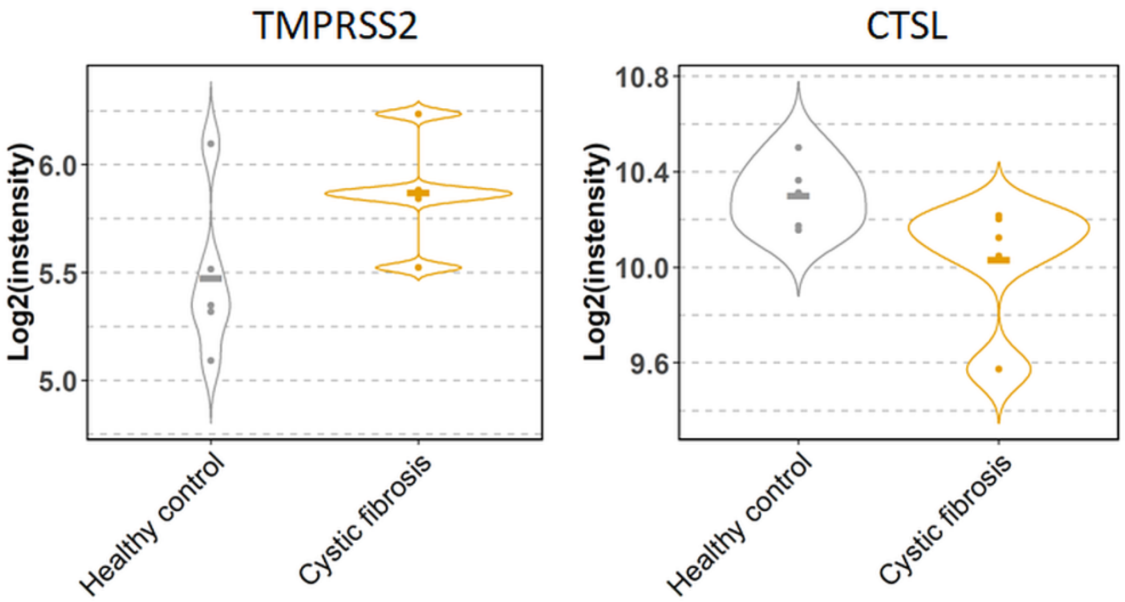


**Supplementary Fig. 7. Expression of TMPRSS2 and cathepsin L (CTSL) in airway epithelial cells of healthy controls (n=5) and patients with cystic fibrosis (n=5) in dataset GSE40445.** No significant difference has been observed between the two groups.

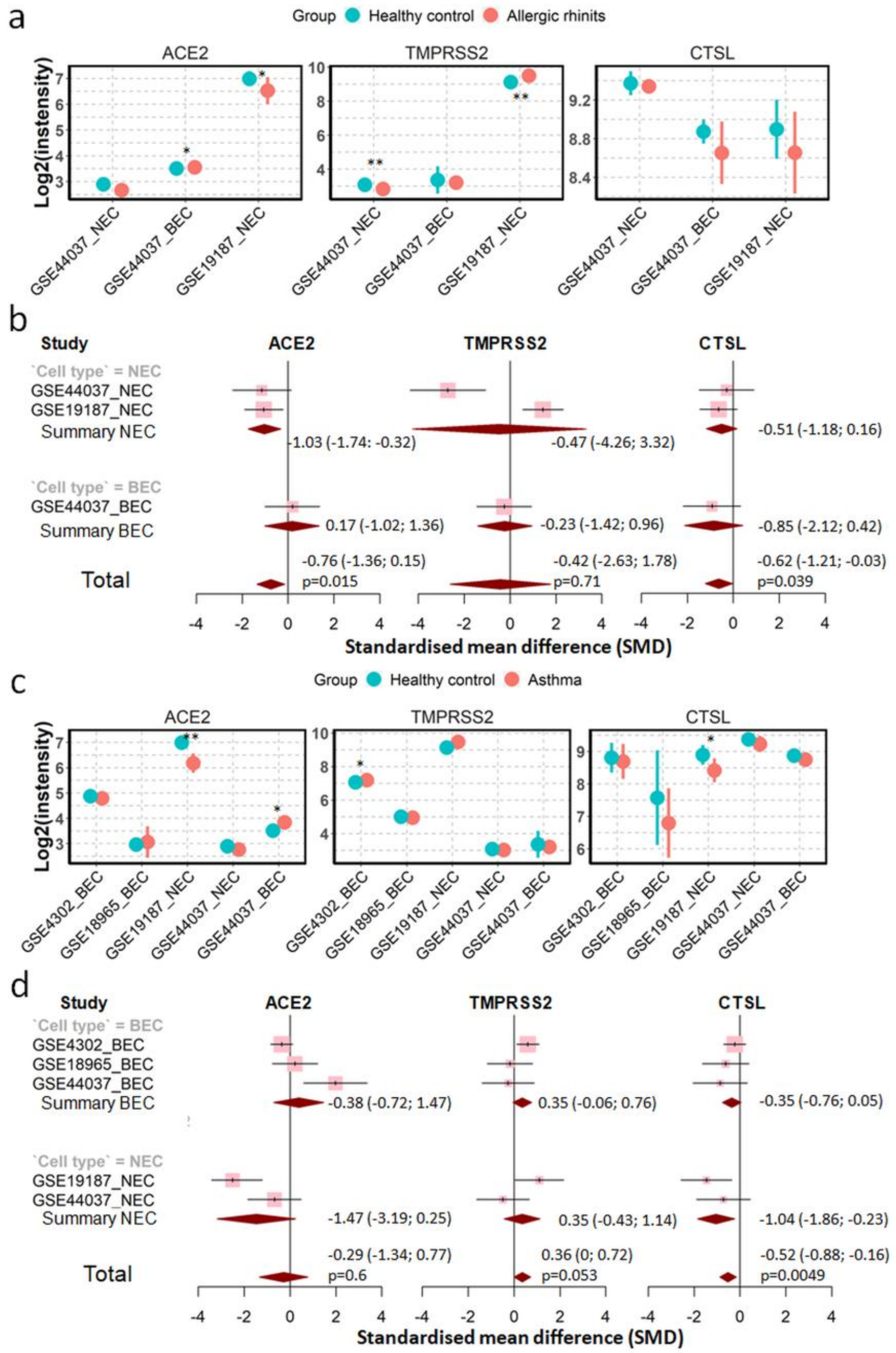


**Supplementary Fig. 8. Expression of ACE2, TMPRSS2 and cathepsin L (CTSL) in airway epithelial cells of healthy controls, patients with asthma and patients with allergic rhinitis. a)** Plot of expression levels of ACE2, TMPRSS2 and CTSL in 3 datasets which contain healthy control and patients with allergic rhinitis. Statistical difference was calculated by Student' t test. *p<0.05, **p<0.01, ***p<0.001. **b)** Forest plot of 3 datasets examining expression of ACE2, TMPRSS2 and CTSL in healthy controls and patients with allergic rhinitis. The x-axis indicates the standardized mean difference (SMD), while the y-axis shows GEO datasets and cell types. The SMD, 95% CI and p values of meta-analysis are depicted. **c)** Plot of expression levels of ACE2, TMPRSS2 and CTSL in 5 datasets which contain healthy control and patients with asthma. **d)** Forest plot of 5 datasets examining expression of ACE2, TMPRSS2 and CTSL in healthy controls and patients with asthma. BEC, bronchial epithelial cells; NEC, nasal epithelial cells.

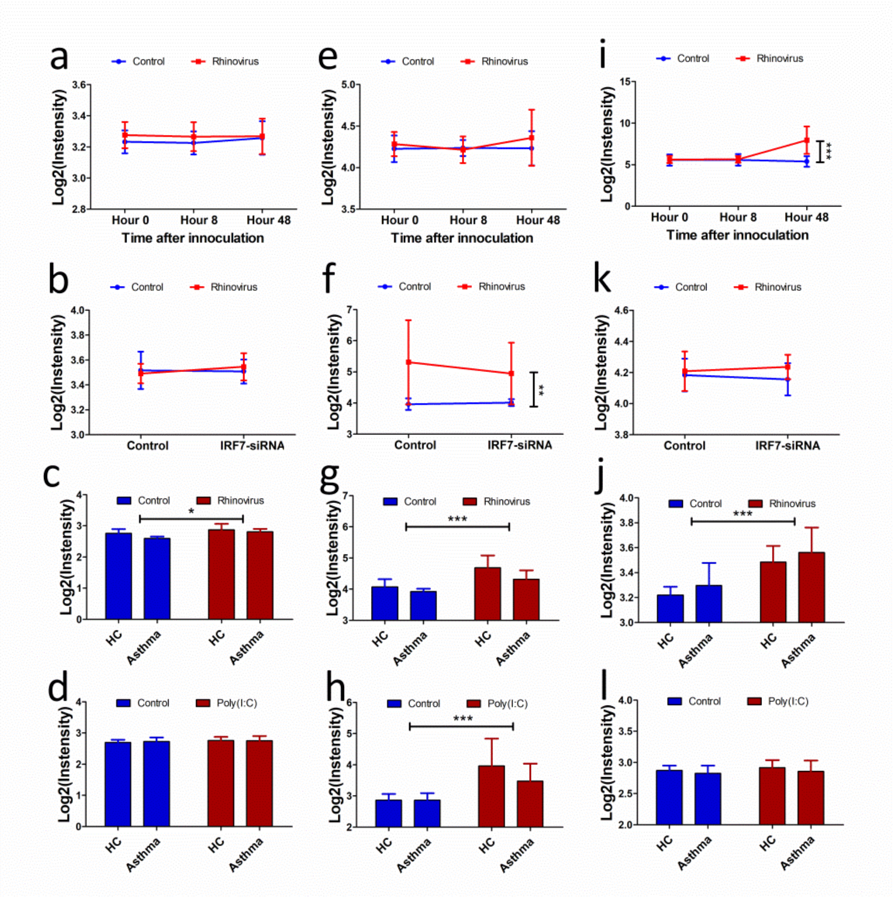


**Supplementary Fig. 9. Expression of IFN-α, IFN-β and IFN-γ in airway epithelial cells after rhinoviral infection or TLR3 activation.** Kinetics of expression of IFN-α (a), IFN-β (e), and IFN-γ (i) in airway epithelial cells of healthy subjects artificially infected with rhinovirus or saline control (data from GSE11348). Expression of IFN-α (b), IFN-β (f), and IFN-γ (j) in airway epithelial cells isolated from healthy subjected and stimulated in vitro with or without rhinovirus in presence or absence of IRF-siRNA (data from GSE70190). Expression of IFN-α (c), IFN-β (g), and IFN-γ (k) in airway epithelial cells isolated from healthy subjected or patients with asthma and stimulated in vitro with rhinovirus or saline control (data from GSE13396). Expression of IFN-α (d), IFN-β (h), and IFN-γ (l) in airway epithelial cells isolated from healthy subjected or patients with asthma and stimulated in vitro with polyI:C or or saline control (data from GSE13396). Statistical analysis was performed using two-way ANOVA, Tukey's test for post hoc analysis was performed after two-way ANOVA analysis. *, $p<0.05$, **, $p<0.01$, ***, $p<0.001$. HC, healthy control; NS, not significant.
